## Supporting Information contains Tables S1-S5 and Figures S1-S3 for "Pharmacokinetic-Pharmacodynamic Trade-offs in SARS-CoV-2 Main Protease Inhibitors Unveiled through Machine Learning and Molecular Dynamics Simulations"

### Supplementary text

#### Summary of parameters and hyperparameters of the Support Vector Machine (model 9)

```
MODEL SUMMARY
=============

Model Name: Model 9 (SVM)
Kernel: rbf
Number of K-Fold Splits: 5
Random State (KF): 37
Test Size: 0.2
Random State (Split): 21

MODEL PARAMETERS
================

Positive class: active
Classes: ['active' 'inactive']
Class weights: None
Intercept: [0.86278088]
Number of support vectors per class: [357 412]
Total number of support vectors: 769
Kernel type: rbf
Degree (for polynomial kernel): 3
Gamma (kernel coefficient): 0.02829116723170816
Coef0 (independent term in kernel): 0.0
Probability estimates: True
Shrinking heuristic: True
Tolerance for stopping criterion: 0.001
Cache size: 200
Maximum iterations: -1
Decision function shape: ovr
Break ties: False

SUPPORT VECTORS
===============

Number of support vectors: 769
Support vector indices:
[   2    6    9   11   22   24   28   34   38   41   42   44   48   49
   52   53   56   58   61   62   64   65   68   75   80   82   85   86
   91   95   98  101  105  107  110  123  132  136  137  139  140  143
  147  148  150  152  153  156  159  160  161  165  167  168  170  171
  172  173  176  179  182  183  184  185  187  189  191  192  194  199
  204  205  208  209  212  218  222  225  238  240  241  243  250  252
  258  259  264  270  276  281  282  286  287  289  290  296  297  301
  303  309  321  326  334  335  336  344  346  348  351  357  364  373
  376  381  382  388  390  394  395  399  403  405  406  407  409  411
  414  416  419  423  426  428  429  430  439  440  444  446  448  450
  453  454  456  457  458  463  471  476  478  487  489  492  502  507
  510  514  517  520  524  526  536  541  542  546  553  554  555  556
  557  560  565  566  567  570  575  576  577  578  590  605  606  621
  622  623  628  635  641  642  644  646  648  652  656  659  661  665
  667  674  676  680  682  684  685  691  698  700  704  709  710  713
```

```
  715  721  726  730  733  734  736  739  747  752  753  760  763  772
  774  779  784  785  787  790  795  796  798  804  805  806  807  810
  814  824  828  829  832  835  838  844  846  852  857  859  860  862
  867  873  879  883  885  887  894  904  915  923  927  933  938  940
  941  952  960  963  965  967  973  977  983  984  985  989  990  992
  995 1006 1010 1011 1015 1016 1019 1021 1023 1027 1032 1038 1041 1042
 1044 1047 1049 1053 1055 1057 1066 1068 1073 1082 1085 1086 1092 1101
 1103 1106 1108 1112 1113 1114 1116 1120 1125 1128 1134 1139 1140 1151
 1153 1154 1155 1161 1162 1166 1174 1179 1181 1184 1188 1189 1192 1193
 1195 1196 1199 1200 1208 1212 1218 1220 1221 1224 1225 1226 1228 1244
 1253 1254 1257 1268 1269 1273 1274   14   16   17   21   23   25   29
   30   31   39   45   46   51   63   70   72   74   77   81   83   87
   96   97  103  106  109  111  112  115  117  121  127  130  133  134
  138  144  145  146  155  162  169  175  181  186  193  196  197  198
  201  213  216  217  221  223  224  228  233  236  237  239  246  247
  253  260  262  263  267  268  269  271  275  279  283  284  291  292
  293  294  298  299  300  302  304  305  308  312  314  318  319  320
  322  323  324  325  327  328  329  332  333  338  339  340  341  345
  350  353  355  356  360  363  365  366  367  368  379  383  385  386
  387  389  396  397  398  401  402  410  412  421  425  433  436  437
  441  449  451  452  455  460  464  466  467  468  469  475  480  481
  482  483  484  485  488  490  493  495  496  498  504  505  506  513
  515  516  519  522  525  527  529  531  533  537  538  539  543  544
  548  549  550  551  561  562  568  573  579  581  583  584  585  586
  592  593  595  596  598  599  610  611  612  613  615  617  619  630
  634  637  640  645  647  649  650  653  654  655  657  658  662  671
  675  688  692  693  694  699  702  705  706  708  714  716  717  718
  719  723  724  725  727  731  740  746  748  749  754  755  756  761
  762  767  768  771  775  776  780  781  782  783  786  788  793  797
  802  808  812  813  815  818  819  821  823  825  827  830  833  843
  849  850  851  853  855  856  861  864  865  872  874  876  884  886
  888  889  891  892  897  902  903  906  907  908  909  910  929  930
  931  932  934  939  943  944  947  950  953  955  957  961  964  969
  970  971  972  979  980  982  987  988  991  994  997  998  999 1008
 1013 1014 1018 1024 1025 1026 1029 1035 1036 1039 1040 1043 1048 1050
 1051 1059 1064 1067 1070 1072 1075 1076 1077 1079 1084 1088 1090 1095
 1096 1097 1098 1100 1105 1109 1117 1118 1119 1122 1123 1127 1129 1131
 1133 1135 1141 1144 1148 1149 1150 1156 1163 1170 1172 1173 1175 1176
 1177 1178 1183 1185 1187 1206 1207 1210 1213 1214 1219 1227 1234 1236
 1238 1239 1243 1245 1246 1250 1251 1258 1261 1265 1266 1270 1271]
Support vectors shape: (769, 36)

SCALER PARAMETERS
=================

Mean: [ 1.28456988e+01  1.11083762e-01 -1.18221415e+00  6.05752648e-01
  1.49235878e+01  4.07652103e+02  2.57534095e-01 -4.34510031e-01
  1.16447679e+00  3.00252872e+01  9.99302489e+00  2.28411591e+00
 -2.25395734e+00  2.30223232e+00 -2.35372658e+00  6.60536099e+00
 -7.61698288e-02  2.97449637e+00  1.72962610e+00 -3.01011603e+00
  2.70663390e+13  3.65201905e+01  3.05128538e+01  7.09602571e+01
  3.31052145e+01  8.06723386e+01  2.62066357e+01  1.19352863e+01
  3.45182939e+00  3.25650377e-01  9.25901267e-01  5.66666120e-04
  8.36178581e-01  1.08653030e+00  5.74255544e+03  3.00766457e-01]
```

```
Scale: [1.44544092e+00 1.15081938e-01 1.38569345e+00 1.94226585e-01
 4.60937248e+00 1.05590021e+02 5.01626151e-02 1.15123692e-01
 1.37684773e-01 1.03973545e+01 1.60289068e-01 1.35423178e-01
 1.22602269e-01 9.67695891e-02 1.58508481e-01 8.04332064e-01
 1.77966012e-01 3.81150605e-01 3.30783452e-01 1.00438939e+00
 8.38390329e+14 1.64850409e+01 1.24323480e+01 2.51408370e+01
 1.83384348e+01 3.76323224e+01 1.38573995e+01 4.80298019e+00
 1.48671832e+00 1.59199674e-01 5.74079732e-02 6.53550481e-04
 8.94009435e-02 3.00379996e-01 3.13034066e+03 1.26700292e-01]
Number of features seen: 36
Feature names: ['MaxAbsEStateIndex' 'MinAbsEStateIndex' 'MinEStateIndex' 'qed' 'SPS'
 'MolWt' 'MaxPartialCharge' 'MinPartialCharge' 'FpDensityMorgan1'
 'BCUT2D_MWHI' 'BCUT2D_MWLOW' 'BCUT2D_CHGHI' 'BCUT2D_CHGLO'
 'BCUT2D_LOGPHI' 'BCUT2D_LOGPLOW' 'BCUT2D_MRHI' 'BCUT2D_MRLOW' 'AvgIpc'
 'BalabanJ' 'HallKierAlpha' 'Ipc' 'PEOE_VSA7' 'SMR_VSA10' 'SMR_VSA7'
 'SlogP_VSA2' 'TPSA' 'VSA_EState2' 'VSA_EState6' 'MolLogP'
 'CalcAsphericity' 'CalcEccentricity' 'CalcInertialShapeFactor' 'CalcNPR2'
 'CalcPBF' 'CalcPMI2' 'CalcSpherocityIndex']

DATA INFORMATION
================

Number of Descriptors: 36
Descriptors Used: MaxAbsEStateIndex, MinAbsEStateIndex, MinEStateIndex, qed, SPS, MolWt,
MaxPartialCharge, MinPartialCharge, FpDensityMorgan1, BCUT2D_MWHI, BCUT2D_MWLOW, BCUT2D_CHGHI,
BCUT2D_CHGLO, BCUT2D_LOGPHI, BCUT2D_LOGPLOW, BCUT2D_MRHI, BCUT2D_MRLOW, AvgIpc, BalabanJ,
HallKierAlpha, Ipc, PEOE_VSA7, SMR_VSA10, SMR_VSA7, SlogP_VSA2, TPSA, VSA_EState2,
VSA_EState6, MolLogP, CalcAsphericity, CalcEccentricity, CalcInertialShapeFactor,
CalcNPR2, CalcPBF, CalcPMI2, CalcSpherocityIndex

SAMPLE SIZES
============

Train Set Size: 1276 compounds
Test Set Size: 319 compounds
Total Size: 1595 compounds

CLASS DISTRIBUTION
==================

Train Set:
- Active: 475 compounds
- Inactive: 801 compounds

Test Set:
- Active: 119 compounds
- Inactive: 200 compounds

PERFORMANCE METRICS
===================

Train Set:
- Accuracy: 0.8393
- Precision: 0.8111
```

```
- Recall: 0.7411
- F1 Score: 0.7745
- ROC AUC: 0.9052

Test Set:
- Accuracy: 0.7931
- Precision: 0.7524
- Recall: 0.6639
- F1 Score: 0.7054
- ROC AUC: 0.8556

CONFUSION MATRIX (Train)
=======================
[[352 123]
 [ 82 719]]

CONFUSION MATRIX (Test)
======================
[[ 79  40]
 [ 26 174]]
```

### Summary of parameters and hyperparameters of the Logistic Regression (model 10)

```
MODEL SUMMARY
=============

Model Name: Logistic Regression (Model 10)
Solver: newton-cg
Number of K-Fold Splits: 5
Random State (KF): 11
Test Size: 0.25
Random State (Split): 75

MODEL PARAMETERS
================

Positive class: active
Classes: ['active' 'inactive']
Class weights: None
Intercept: [0.84751597]
Maximum iterations: 1000
Actual iterations: 8
Penalty: l2
Tolerance: 0.0001
Multi-class: deprecated
Warm start: False
Fit intercept: True
Dual formulation: False
Regularization strength (C): 1.0
Intercept scaling: 1
Verbose: 0

MODEL COEFFICIENTS (TRANSFORMED DATA - SCALED)
==============================================

Coefficients for class 'active':
MaxAbsEStateIndex: -0.045758
MaxEStateIndex: -0.045758
MinAbsEStateIndex: 0.261263
MinEStateIndex: 0.052071
qed: 0.368401
SPS: -0.135354
MolWt: 0.350470
HeavyAtomMolWt: 0.261952
ExactMolWt: 0.364980
MaxPartialCharge: -0.107447
MinPartialCharge: -0.019285
MaxAbsPartialCharge: 0.033589
MinAbsPartialCharge: -0.151437
FpDensityMorgan1: 0.272892
FpDensityMorgan2: -0.484423
FpDensityMorgan3: 0.182740
BCUT2D_MWHI: 0.028559
BCUT2D_MWLOW: 0.326069
BCUT2D_CHGHI: -0.057932
BCUT2D_CHGLO: -0.391952
```

BCUT2D_LOGPHI: 0.233287
BCUT2D_LOGPLOW: -0.066043
BCUT2D_MRHI: 0.015984
BCUT2D_MRLOW: 0.159191
AvgIpc: 0.177490
BalabanJ: -0.017653
BertzCT: 0.440002
Chi0: -0.022032
Chi0n: -0.089333
Chi0v: -0.184038
Chi1: -0.479127
Chi1n: -0.054224
Chi1v: 0.198553
Chi2n: -1.157726
Chi2v: -0.749092
Chi3n: 0.083263
Chi3v: 0.462206
Chi4n: 0.029118
Chi4v: 0.208155
HallKierAlpha: -0.419543
Ipc: 0.048477
Kappa1: 0.010572
Kappa2: -0.449544
Kappa3: 0.528431
LabuteASA: -0.767676
PEOE_VSA7: 0.242436
SMR_VSA10: -0.875387
SMR_VSA7: 0.539763
SlogP_VSA2: -0.553164
SlogP_VSA6: -0.156374
TPSA: -0.000156
VSA_EState2: -0.513693
VSA_EState6: -0.375356
Phi: 0.208953
MolLogP: -0.600020
MolMR: -0.163335
CalcAsphericity: 0.300836
CalcChi0n: 0.859665
CalcChi0v: -0.462352
CalcChi1n: 0.676144
CalcChi1v: -0.109783
CalcChi2n: -0.591301
CalcChi2v: -0.651634
CalcChi3n: -0.162703
CalcChi3v: -0.261868
CalcChi4n: 0.901846
CalcChi4v: 0.343544
CalcEccentricity: -0.614794
CalcExactMolWt: 0.364980
CalcHallKierAlpha: -0.419543
CalcInertialShapeFactor: 0.114054
CalcKappa1: 0.180560
CalcKappa2: -0.449544
CalcKappa3: 0.528431

```
CalcLabuteASA: -0.334686
CalcNPR1: -0.791124
CalcNPR2: -0.149366
CalcPBF: -0.371305
CalcPMI1: 0.729673
CalcPMI2: 0.278562
CalcPMI3: 1.143904
CalcPhi: -0.136720
CalcRadiusOfGyration: -0.691894
CalcSpherocityIndex: 0.484147
CalcTPSA: -0.000156

MODEL COEFFICIENTS (ORIGINAL DATA - UNSCALED)
============================================

Coefficients for class 'active':
MaxAbsEStateIndex: -0.031657
MaxEStateIndex: -0.031657
MinAbsEStateIndex: 2.270231
MinEStateIndex: 0.037577
qed: 1.896756
SPS: -0.029365
MolWt: 0.003319
HeavyAtomMolWt: 0.002646
ExactMolWt: 0.003458
MaxPartialCharge: -2.141974
MinPartialCharge: -0.167517
MaxAbsPartialCharge: 0.294682
MinAbsPartialCharge: -3.276943
FpDensityMorgan1: 1.982005
FpDensityMorgan2: -2.687581
FpDensityMorgan3: 0.743010
BCUT2D_MWHI: 0.002747
BCUT2D_MWLOW: 2.034253
BCUT2D_CHGHI: -0.427782
BCUT2D_CHGLO: -3.196941
BCUT2D_LOGPHI: 2.410748
BCUT2D_LOGPLOW: -0.416651
BCUT2D_MRHI: 0.019872
BCUT2D_MRLOW: 0.894504
AvgIpc: 0.465669
BalabanJ: -0.053367
BertzCT: 0.001345
Chi0: -0.003949
Chi0n: -0.019977
Chi0v: -0.041809
Chi1: -0.130236
Chi1n: -0.020580
Chi1v: 0.072680
Chi2n: -0.507627
Chi2v: -0.304748
Chi3n: 0.050876
Chi3v: 0.247874
Chi4n: 0.023668
```

```
Chi4v: 0.143030
HallKierAlpha: -0.417710
Ipc: 0.000000
Kappa1: 0.001779
Kappa2: -0.162286
Kappa3: 0.288044
LabuteASA: -0.017484
PEOE_VSA7: 0.014706
SMR_VSA10: -0.070412
SMR_VSA7: 0.021470
SlogP_VSA2: -0.030164
SlogP_VSA6: -0.007691
TPSA: -0.000004
VSA_EState2: -0.037070
VSA_EState6: -0.078151
Phi: 0.094010
MolLogP: -0.403587
MolMR: -0.005881
CalcAsphericity: 1.889679
CalcChi0n: 0.075879
CalcChi0v: -0.126172
CalcChi1n: 0.118212
CalcChi1v: -0.052301
CalcChi2n: -0.409063
CalcChi2v: -0.398396
CalcChi3n: -0.165065
CalcChi3v: -0.228856
CalcChi4n: 1.346273
CalcChi4v: 0.424265
CalcEccentricity: -10.709207
CalcExactMolWt: 0.003458
CalcHallKierAlpha: -0.417710
CalcInertialShapeFactor: 174.514104
CalcKappa1: 0.098103
CalcKappa2: -0.162286
CalcKappa3: 0.288044
CalcLabuteASA: -0.006206
CalcNPR1: -5.970694
CalcNPR2: -1.670741
CalcPBF: -1.236118
CalcPMI1: 0.000413
CalcPMI2: 0.000089
CalcPMI3: 0.000312
CalcPhi: -0.223779
CalcRadiusOfGyration: -1.292396
CalcSpherocityIndex: 3.821201
CalcTPSA: -0.000004

Intercept (adjusted for original scale): -11.559802

SCALER PARAMETERS
=================

Mean: [ 1.28456988e+01  1.28456988e+01  1.11083762e-01 -1.18221415e+00
```

```
 6.05752648e-01  1.49235878e+01  4.07652103e+02  3.87171060e+02
 4.07059810e+02  2.57534095e-01 -4.34510031e-01  4.36304609e-01
 2.55739517e-01  1.16447679e+00  1.95428086e+00  2.69015918e+00
 3.00252872e+01  9.99302489e+00  2.28411591e+00 -2.25395734e+00
 2.30223232e+00 -2.35372658e+00  6.60536099e+00 -7.61698288e-02
 2.97449637e+00  1.72962610e+00  1.06468946e+03  2.03704146e+01
 1.58108119e+01  1.65420058e+01  1.38596590e+01  9.20962739e+00
 9.85161933e+00  7.02112286e+00  7.75012797e+00  4.87873394e+00
 5.44639202e+00  3.39717158e+00  3.86481627e+00 -3.01011603e+00
 2.70663390e+13  1.95014745e+01  8.18041191e+00  4.22907615e+00
 1.70332192e+02  3.65201905e+01  3.05128538e+01  7.09602571e+01
 3.31052145e+01  5.60508896e+01  8.06723386e+01  2.62066357e+01
 1.19352863e+01  5.58033418e+00  3.45182939e+00  1.09557148e+02
 3.25650377e-01  3.41267468e+01  1.45393965e+01  1.75112466e+01
 8.02236447e+00  5.09258419e+00  5.70838880e+00  3.27900968e+00
 3.70754978e+00  2.08844072e+00  2.40594597e+00  9.25901267e-01
 4.07059810e+02 -3.01011603e+00  5.66666120e-04  6.73984888e+00
 8.18041191e+00  4.22907615e+00  1.98952119e+02  3.49076820e-01
 8.36178581e-01  1.08653030e+00  2.49356433e+03  5.74255544e+03
 6.89745992e+03  1.90452974e+00  4.16332835e+00  3.00766457e-01
 8.06723386e+01]
Scale: [1.44544092e+00 1.44544092e+00 1.15081938e-01 1.38569345e+00
 1.94226585e-01 4.60937248e+00 1.05590021e+02 9.89845558e+01
 1.05549397e+02 5.01626151e-02 1.15123692e-01 1.13984631e-01
 4.62127792e-02 1.37684773e-01 1.80244833e-01 2.45945269e-01
 1.03973545e+01 1.60289068e-01 1.35423178e-01 1.22602269e-01
 9.67695891e-02 1.58508481e-01 8.04332064e-01 1.77966012e-01
 3.81150605e-01 3.30783452e-01 3.27242193e+02 5.57899108e+00
 4.47183281e+00 4.40189648e+00 3.67890752e+00 2.63471781e+00
 2.73187461e+00 2.28066280e+00 2.45807363e+00 1.63659560e+00
 1.86468215e+00 1.23026759e+00 1.45531880e+00 1.00438939e+00
 8.38390329e+14 5.94286298e+00 2.77008161e+00 1.83454600e+00
 4.39073747e+01 1.64850409e+01 1.24323480e+01 2.51408370e+01
 1.83384348e+01 2.03313352e+01 3.76323224e+01 1.38573995e+01
 4.80298019e+00 2.22266656e+00 1.48671832e+00 2.77710143e+01
 1.59199674e-01 1.13293876e+01 3.66447066e+00 5.71974032e+00
 2.09906609e+00 1.44550056e+00 1.63564554e+00 9.85687926e-01
 1.14424529e+00 6.69883294e-01 8.09740274e-01 5.74079732e-02
 1.05549397e+02 1.00438939e+00 6.53550481e-04 1.84051746e+00
 2.77008161e+00 1.83454600e+00 5.39335335e+01 1.32501103e-01
 8.94009435e-02 3.00379996e-01 1.76518758e+03 3.13034066e+03
 3.66674055e+03 6.10961628e-01 5.35357266e-01 1.26700292e-01
 3.76323224e+01]
Number of features seen: 85
Feature names: ['MaxAbsEStateIndex' 'MaxEStateIndex' 'MinAbsEStateIndex' 'MinEStateIndex'
 'qed' 'SPS' 'MolWt' 'HeavyAtomMolWt' 'ExactMolWt' 'MaxPartialCharge'
 'MinPartialCharge' 'MaxAbsPartialCharge' 'MinAbsPartialCharge'
 'FpDensityMorgan1' 'FpDensityMorgan2' 'FpDensityMorgan3' 'BCUT2D_MWHI'
 'BCUT2D_MWLOW' 'BCUT2D_CHGHI' 'BCUT2D_CHGLO' 'BCUT2D_LOGPHI'
 'BCUT2D_LOGPLOW' 'BCUT2D_MRHI' 'BCUT2D_MRLOW' 'AvgIpc' 'BalabanJ'
 'BertzCT' 'Chi0' 'Chi0n' 'Chi0v' 'Chi1' 'Chi1n' 'Chi1v' 'Chi2n' 'Chi2v'
 'Chi3n' 'Chi3v' 'Chi4n' 'Chi4v' 'HallKierAlpha' 'Ipc' 'Kappa1' 'Kappa2'
 'Kappa3' 'LabuteASA' 'PEOE_VSA7' 'SMR_VSA10' 'SMR_VSA7' 'SlogP_VSA2'
 'SlogP_VSA6' 'TPSA' 'VSA_EState2' 'VSA_EState6' 'Phi' 'MolLogP' 'MolMR'
```

```
 'CalcAsphericity' 'CalcChi0n' 'CalcChi0v' 'CalcChi1n' 'CalcChi1v'
 'CalcChi2n' 'CalcChi2v' 'CalcChi3n' 'CalcChi3v' 'CalcChi4n' 'CalcChi4v'
 'CalcEccentricity' 'CalcExactMolWt' 'CalcHallKierAlpha'
 'CalcInertialShapeFactor' 'CalcKappa1' 'CalcKappa2' 'CalcKappa3'
 'CalcLabuteASA' 'CalcNPR1' 'CalcNPR2' 'CalcPBF' 'CalcPMI1' 'CalcPMI2'
 'CalcPMI3' 'CalcPhi' 'CalcRadiusOfGyration' 'CalcSpherocityIndex'
 'CalcTPSA']

DATA INFORMATION
================

Number of Descriptors: 85
Descriptors Used: MaxAbsEStateIndex, MaxEStateIndex, MinAbsEStateIndex, MinEStateIndex, qed,
SPS, MolWt, HeavyAtomMolWt, ExactMolWt, MaxPartialCharge, MinPartialCharge, MaxAbsPartialCharge,
MinAbsPartialCharge, FpDensityMorgan1, FpDensityMorgan2, FpDensityMorgan3, BCUT2D_MWHI,
BCUT2D_MWLOW, BCUT2D_CHGHI, BCUT2D_CHGLO, BCUT2D_LOGPHI, BCUT2D_LOGPLOW, BCUT2D_MRHI,
BCUT2D_MRLOW, AvgIpc, BalabanJ, BertzCT, Chi0, Chi0n, Chi0v, Chi1, Chi1n, Chi1v, Chi2n, Chi2v,
Chi3n, Chi3v, Chi4n, Chi4v, HallKierAlpha, Ipc, Kappa1, Kappa2, Kappa3, LabuteASA, PEOE_VSA7,
SMR_VSA10, SMR_VSA7, SlogP_VSA2, SlogP_VSA6, TPSA, VSA_EState2, VSA_EState6, Phi, MolLogP,
MolMR, CalcAsphericity, CalcChi0n, CalcChi0v, CalcChi1n, CalcChi1v, CalcChi2n, CalcChi2v,
CalcChi3n, CalcChi3v, CalcChi4n, CalcChi4v, CalcEccentricity, CalcExactMolWt, CalcHallKierAlpha,
CalcInertialShapeFactor, CalcKappa1, CalcKappa2, CalcKappa3, CalcLabuteASA, CalcNPR1, CalcNPR2,
CalcPBF, CalcPMI1, CalcPMI2, CalcPMI3, CalcPhi, CalcRadiusOfGyration, CalcSpherocityIndex, CalcTPSA

SAMPLE SIZES
============

Train Set Size: 1196 compounds
Test Set Size: 399 compounds
Total Size: 1595 compounds

CLASS DISTRIBUTION
==================

Train Set:
- Active: 445 compounds
- Inactive: 751 compounds

Test Set:
- Active: 149 compounds
- Inactive: 250 compounds

PERFORMANCE METRICS
===================

Train Set:
- Accuracy: 0.7809
- Precision: 0.7259
- Recall: 0.6607
- F1 Score: 0.6918
- ROC AUC: 0.8481

Test Set:
- Accuracy: 0.7619
```

```
- Precision: 0.7143
- Recall: 0.6040
- F1 Score: 0.6545
- ROC AUC: 0.8270

CONFUSION MATRIX (Train)
=======================
[[294 151]
 [111 640]]

CONFUSION MATRIX (Test)
======================
[[ 90  59]
 [ 36 214]]
```

### Tables

**Table S1. Different conditions in construction of the ML models.** The condition refers to molecular descriptors and dataset with fluorescence data ($N = 1595$), FRET ($N = 353$), and combined data FRET/fluorescence/SPR ($N = 1943$) used in building ML models. Abbreviation: ND = number of descriptors; Dim. = dimension; Thresh. = threshold; Meas. = measurement.

| Cond. | Meas. | Dim. | Thresh. | Selected descriptors | ND |
|---|---|---|---|---|---|
| 1 | FRET | 3D | 0.1 | MaxAbsEStateIndex | 1 |
| 2 | FRET | 3D | 0.3 | MaxAbsEStateIndex, qed, SPS | 3 |
| 3 | FRET | 3D | 0.4 | MaxAbsEStateIndex, MinEStateIndex, qed, SPS, Ipc | 5 |
| 4 | FRET | 2D | 0.5 | MaxAbsEStateIndex, MinEStateIndex, qed, SPS, FpDensityMorgan1, Ipc | 6 |
| 5 | FRET | 3D | 0.5 | MaxAbsEStateIndex, MinEStateIndex, qed, SPS, FpDensityMorgan1, Ipc, CalcAsphericity | 7 |
| 6 | FRET | 2D | 0.6 | MaxAbsEStateIndex, MinEStateIndex, qed, SPS, FpDensityMorgan1, Ipc, MolLogP | 7 |
| 7 | FRET | 3D | 0.6 | MaxAbsEStateIndex, MinEStateIndex, qed, SPS, FpDensityMorgan1, Ipc, MolLogP, CalcAsphericity | 8 |
| 8 | FRET | 2D | 0.7 | MaxAbsEStateIndex, MinAbsEStateIndex, MinEStateIndex, qed, SPS, MolWt, FpDensityMorgan1, Ipc, Kappa3, SMR_VSA10, SMR_VSA7, MolLogP | 12 |
| 9 | FRET | 3D | 0.7 | MaxAbsEStateIndex, MinAbsEStateIndex, MinEStateIndex, qed, SPS, MolWt, FpDensityMorgan1, Ipc, Kappa3, SMR_VSA10, SMR_VSA7, MolLogP, CalcAsphericity, CalcNPR2 | 14 |
| 10 | FRET | 2D | 0.8 | MaxAbsEStateIndex, MinAbsEStateIndex, MinEStateIndex, qed, SPS, MolWt, FpDensityMorgan1, AvgIpc, BalabanJ, Ipc, Kappa3, SMR_VSA10, SMR_VSA7, SlogP_VSA2, MolLogP | 15 |
| 11 | FRET | 3D | 0.8 | MaxAbsEStateIndex, MinAbsEStateIndex, MinEStateIndex, qed, SPS, MolWt, FpDensityMorgan1, AvgIpc, BalabanJ, Ipc, Kappa3, SMR_VSA10, SMR_VSA7, SlogP_VSA2, MolLogP, CalcAsphericity, CalcInertialShapeFactor, CalcNPR2, CalcPBF, CalcSpherocityIndex | 20 |
| 12 | FRET | 2D | 0.9 | MaxAbsEStateIndex, MinAbsEStateIndex, MinEStateIndex, qed, SPS, MolWt, FpDensityMorgan1, AvgIpc, BalabanJ, BertzCT, HallKierAlpha, Ipc, Kappa3, SMR_VSA10, SMR_VSA7, SlogP_VSA2, TPSA, NumHAcceptors, RingCount, MolLogP | 20 |
| 13 | FRET | 3D | 0.9 | MaxAbsEStateIndex, MinAbsEStateIndex, MinEStateIndex, qed, SPS, MolWt, FpDensityMorgan1, AvgIpc, BalabanJ, BertzCT, HallKierAlpha, Ipc, Kappa3, SMR_VSA10, SMR_VSA7, SlogP_VSA2, TPSA, NumHAcceptors, RingCount, MolLogP, CalcAsphericity, CalcEccentricity, CalcInertialShapeFactor, CalcKappa1, CalcNPR2, CalcPBF, CalcPMI2, CalcRadiusOfGyration, CalcSpherocityIndex | 29 |
| 14 | FRET | 2D | 1 | MaxAbsEStateIndex, MaxEStateIndex, MinAbsEStateIndex, MinEStateIndex, qed, SPS, MolWt, HeavyAtomMolWt, ExactMolWt, NumValenceElectrons, FpDensityMorgan1, FpDensityMorgan2, FpDensityMorgan3, AvgIpc, BalabanJ, BertzCT, Chi0, Chi0n, Chi0v, Chi1, Chi1n, Chi1v, Chi2n, Chi2v, Chi3n, Chi3v, Chi4n, Chi4v, HallKierAlpha, Ipc, Kappa1, Kappa2, Kappa3, LabuteASA, SMR_VSA10, SMR_VSA7, SlogP_VSA2, SlogP_VSA6, TPSA, HeavyAtomCount, NOCount, NumHAcceptors, NumHeteroatoms, Phi, RingCount, MolLogP, MolMR | 47 |

Continued on next page

| Cond. | Meas. | Dim. | Thresh. | Selected descriptors | ND |
|---|---|---|---|---|---|
| 15 | FRET | 3D | 1 | MaxAbsEStateIndex, MaxEStateIndex, MinAbsEStateIndex, MinEStateIndex, qed, SPS, MolWt, HeavyAtomMolWt, ExactMolWt, NumValenceElectrons, FpDensityMorgan1, FpDensityMorgan2, FpDensityMorgan3, AvgIpc, BalabanJ, BertzCT, Chi0, Chi0n, Chi0v, Chi1, Chi1n, Chi1v, Chi2n, Chi2v, Chi3n, Chi3v, Chi4n, Chi4v, HallKierAlpha, Ipc, Kappa1, Kappa2, Kappa3, LabuteASA, SMR_VSA10, SMR_VSA7, SlogP_VSA2, SlogP_VSA6, TPSA, HeavyAtomCount, NOCount, NumHAcceptors, NumHeteroatoms, Phi, RingCount, MolLogP, MolMR, CalcAsphericity, CalcChi0n, CalcChi0v, CalcChi1n, CalcChi1v, CalcChi2n, CalcChi2v, CalcChi3n, CalcChi3v, CalcChi4n, CalcChi4v, CalcEccentricity, CalcHallKierAlpha, CalcInertialShapeFactor, CalcKappa1, CalcKappa2, CalcKappa3, CalcLabuteASA, CalcNPR1, CalcNPR2, CalcNumAtoms, CalcNumHBA, CalcNumHeavyAtoms, CalcNumHeteroatoms, CalcNumLipinskiHBA, CalcNumRings, CalcNumRotatableBonds, CalcPBF, CalcPMI1, CalcPMI2, CalcPMI3, CalcPhi, CalcRadiusOfGyration, CalcSpherocityIndex, CalcTPSA | 82 |
| 16 | FRET/fluor/SPR | 3D | 0.1 | MaxAbsEStateIndex | 1 |
| 17 | FRET/fluor/SPR | 3D | 0.2 | MaxAbsEStateIndex, Ipc | 2 |
| 18 | FRET/fluor/SPR | 3D | 0.3 | MaxAbsEStateIndex, qed, SPS, Ipc | 4 |
| 19 | FRET/fluor/SPR | 3D | 0.4 | MaxAbsEStateIndex, MinEStateIndex, qed, SPS, Ipc | 5 |
| 20 | FRET/fluor/SPR | 2D | 0.5 | MaxAbsEStateIndex, MinEStateIndex, qed, SPS, FpDensityMorgan1, Ipc | 6 |
| 21 | FRET/fluor/SPR | 3D | 0.5 | MaxAbsEStateIndex, MinEStateIndex, qed, SPS, FpDensityMorgan1, Ipc, CalcAsphericity | 7 |
| 22 | FRET/fluor/SPR | 2D | 0.6 | MaxAbsEStateIndex, MinEStateIndex, qed, SPS, FpDensityMorgan1, BalabanJ, Ipc, MolLogP | 8 |
| 23 | FRET/fluor/SPR | 3D | 0.6 | MaxAbsEStateIndex, MinEStateIndex, qed, SPS, FpDensityMorgan1, BalabanJ, Ipc, MolLogP, CalcAsphericity, CalcNPR2 | 10 |
| 24 | FRET/fluor/SPR | 2D | 0.7 | MaxAbsEStateIndex, MinAbsEStateIndex, MinEStateIndex, qed, SPS, FpDensityMorgan1, AvgIpc, BalabanJ, Ipc, MolLogP | 10 |
| 25 | FRET/fluor/SPR | 3D | 0.7 | MaxAbsEStateIndex, MinAbsEStateIndex, MinEStateIndex, qed, SPS, FpDensityMorgan1, AvgIpc, BalabanJ, Ipc, MolLogP, CalcAsphericity, CalcInertialShapeFactor, CalcNPR2 | 13 |
| 26 | FRET/fluor/SPR | 2D | 0.8 | MaxAbsEStateIndex, MinAbsEStateIndex, MinEStateIndex, qed, SPS, MolWt, FpDensityMorgan1, AvgIpc, BalabanJ, Ipc, SMR_VSA7, SlogP_VSA2, TPSA, RingCount, MolLogP | 15 |
| 27 | FRET/fluor/SPR | 3D | 0.8 | MaxAbsEStateIndex, MinAbsEStateIndex, MinEStateIndex, qed, SPS, MolWt, FpDensityMorgan1, AvgIpc, BalabanJ, Ipc, SMR_VSA7, SlogP_VSA2, TPSA, RingCount, MolLogP, CalcAsphericity, CalcInertialShapeFactor, CalcNPR2, CalcPBF | 19 |
| 28 | FRET/fluor/SPR | 2D | 0.9 | MaxAbsEStateIndex, MinAbsEStateIndex, MinEStateIndex, qed, SPS, MolWt, FpDensityMorgan1, AvgIpc, BalabanJ, BertzCT, HallKierAlpha, Ipc, Kappa3, SMR_VSA7, SlogP_VSA2, TPSA, NumHAcceptors, RingCount, MolLogP | 19 |
| 29 | FRET/fluor/SPR | 3D | 0.9 | MaxAbsEStateIndex, MinAbsEStateIndex, MinEStateIndex, qed, SPS, MolWt, FpDensityMorgan1, AvgIpc, BalabanJ, BertzCT, HallKierAlpha, Ipc, Kappa3, SMR_VSA7, SlogP_VSA2, TPSA, NumHAcceptors, RingCount, MolLogP, CalcAsphericity, CalcEccentricity, CalcInertialShapeFactor, CalcKappa1, CalcNPR2, CalcPBF, CalcPMI2, CalcRadiusOfGyration, CalcSpherocityIndex | 28 |

Continued on next page

| Cond. | Meas. | Dim. | Thresh. | Selected descriptors | ND |
|---|---|---|---|---|---|
| 30 | FRET/fluor/SPR | 2D | 1 | MaxAbsEStateIndex, MaxEStateIndex, MinAbsEStateIndex, MinEStateIndex, qed, SPS, MolWt, HeavyAtomMolWt, ExactMolWt, NumValenceElectrons, FpDensityMorgan1, FpDensityMorgan2, FpDensityMorgan3, AvgIpc, BalabanJ, BertzCT, Chi0, Chi0n, Chi0v, Chi1, Chi1n, Chi1v, Chi2n, Chi2v, Chi3n, Chi3v, Chi4n, Chi4v, HallKierAlpha, Ipc, Kappa1, Kappa2, Kappa3, LabuteASA, SMR_VSA7, SlogP_VSA2, TPSA, HeavyAtomCount, NOCount, NumHAcceptors, NumHeteroatoms, Phi, RingCount, MolLogP, MolMR | 45 |
| 31 | FRET/fluor/SPR | 3D | 1 | MaxAbsEStateIndex, MaxEStateIndex, MinAbsEStateIndex, MinEStateIndex, qed, SPS, MolWt, HeavyAtomMolWt, ExactMolWt, NumValenceElectrons, FpDensityMorgan1, FpDensityMorgan2, FpDensityMorgan3, AvgIpc, BalabanJ, BertzCT, Chi0, Chi0n, Chi0v, Chi1, Chi1n, Chi1v, Chi2n, Chi2v, Chi3n, Chi3v, Chi4n, Chi4v, HallKierAlpha, Ipc, Kappa1, Kappa2, Kappa3, LabuteASA, SMR_VSA7, SlogP_VSA2, TPSA, HeavyAtomCount, NOCount, NumHAcceptors, NumHeteroatoms, Phi, RingCount, MolLogP, MolMR, CalcAsphericity, CalcChi0n, CalcChi0v, CalcChi1n, CalcChi1v, CalcChi2n, CalcChi2v, CalcChi3n, CalcChi3v, CalcChi4n, CalcChi4v, CalcEccentricity, CalcExactMolWt, CalcHallKierAlpha, CalcInertialShapeFactor, CalcKappa1, CalcKappa2, CalcKappa3, CalcLabuteASA, CalcNPR1, CalcNPR2, CalcNumAtoms, CalcNumHBA, CalcNumHeavyAtoms, CalcNumHeteroatoms, CalcNumLipinskiHBA, CalcNumRings, CalcNumRotatableBonds, CalcPBF, CalcPMI1, CalcPMI2, CalcPMI3, CalcPhi, CalcRadiusOfGyration, CalcSpherocityIndex, CalcTPSA | 81 |
| 32 | fluor | 3D | 0.2 | MaxAbsEStateIndex | 1 |
| 33 | fluor | 3D | 0.3 | MaxAbsEStateIndex, SPS, MinPartialCharge, Ipc | 4 |
| 34 | fluor | 3D | 0.4 | MaxAbsEStateIndex, MinEStateIndex, qed, SPS, MaxPartialCharge, MinPartialCharge, BCUT2D_MWHI, Ipc | 8 |
| 35 | fluor | 3D | 0.5 | MaxAbsEStateIndex, MinEStateIndex, qed, SPS, MaxPartialCharge, MinPartialCharge, BCUT2D_MWHI, BalabanJ, Ipc | 9 |
| 36 | fluor | 2D | 0.6 | MaxAbsEStateIndex, MinAbsEStateIndex, MinEStateIndex, qed, SPS, MaxPartialCharge, MinPartialCharge, FpDensityMorgan1, BCUT2D_MWHI, BCUT2D_MWLOW, BCUT2D_MRLOW, BalabanJ, Ipc, PEOE_VSA7, SMR_VSA10, MolLogP | 16 |
| 37 | fluor | 3D | 0.6 | MaxAbsEStateIndex, MinAbsEStateIndex, MinEStateIndex, qed, SPS, MaxPartialCharge, MinPartialCharge, FpDensityMorgan1, BCUT2D_MWHI, BCUT2D_MWLOW, BCUT2D_MRLOW, BalabanJ, Ipc, PEOE_VSA7, SMR_VSA10, MolLogP, CalcAsphericity, CalcInertialShapeFactor, CalcNPR2 | 19 |
| 38 | fluor | 2D | 0.7 | MaxAbsEStateIndex, MinAbsEStateIndex, MinEStateIndex, qed, SPS, MaxPartialCharge, MinPartialCharge, FpDensityMorgan1, BCUT2D_MWHI, BCUT2D_MWLOW, BCUT2D_MRHI, BCUT2D_MRLOW, AvgIpc, BalabanJ, Ipc, PEOE_VSA7, SMR_VSA10, MolLogP | 18 |
| 39 | fluor | 3D | 0.7 | MaxAbsEStateIndex, MinAbsEStateIndex, MinEStateIndex, qed, SPS, MaxPartialCharge, MinPartialCharge, FpDensityMorgan1, BCUT2D_MWHI, BCUT2D_MWLOW, BCUT2D_MRHI, BCUT2D_MRLOW, AvgIpc, BalabanJ, Ipc, PEOE_VSA7, SMR_VSA10, MolLogP, CalcAsphericity, CalcInertialShapeFactor, CalcNPR2 | 21 |

Continued on next page

| Cond. | Meas. | Dim. | Thresh. | Selected descriptors | ND |
|---|---|---|---|---|---|
| 40 | fluor | 2D | 0.8 | MaxAbsEStateIndex, MinAbsEStateIndex, MinEStateIndex, qed, SPS, MaxPartialCharge, MinPartialCharge, FpDensityMorgan1, BCUT2D_MWHI, BCUT2D_MWLOW, BCUT2D_CHGHI, BCUT2D_LOGPHI, BCUT2D_MRHI, BCUT2D_MRLOW, AvgIpc, BalabanJ, Ipc, PEOE_VSA7, SMR_VSA10, SlogP_VSA2, TPSA, VSA_EState2, MolLogP | 23 |
| 41 | fluor | 3D | 0.8 | MaxAbsEStateIndex, MinAbsEStateIndex, MinEStateIndex, qed, SPS, MaxPartialCharge, MinPartialCharge, FpDensityMorgan1, BCUT2D_MWHI, BCUT2D_MWLOW, BCUT2D_CHGHI, BCUT2D_LOGPHI, BCUT2D_MRHI, BCUT2D_MRLOW, AvgIpc, BalabanJ, Ipc, PEOE_VSA7, SMR_VSA10, SlogP_VSA2, TPSA, VSA_EState2, MolLogP, CalcAsphericity, CalcInertialShapeFactor, CalcNPR2, CalcPBF | 27 |
| 42 | fluor | 2D | 0.9 | MaxAbsEStateIndex, MinAbsEStateIndex, MinEStateIndex, qed, SPS, MolWt, MaxPartialCharge, MinPartialCharge, FpDensityMorgan1, BCUT2D_MWHI, BCUT2D_MWLOW, BCUT2D_CHGHI, BCUT2D_CHGLO, BCUT2D_LOGPHI, BCUT2D_LOGPLOW, BCUT2D_MRHI, BCUT2D_MRLOW, AvgIpc, BalabanJ, HallKierAlpha, Ipc, PEOE_VSA7, SMR_VSA10, SMR_VSA7, SlogP_VSA2, TPSA, VSA_EState2, VSA_EState6, MolLogP | 29 |
| 43 | fluor | 3D | 0.9 | MaxAbsEStateIndex, MinAbsEStateIndex, MinEStateIndex, qed, SPS, MolWt, MaxPartialCharge, MinPartialCharge, FpDensityMorgan1, BCUT2D_MWHI, BCUT2D_MWLOW, BCUT2D_CHGHI, BCUT2D_CHGLO, BCUT2D_LOGPHI, BCUT2D_LOGPLOW, BCUT2D_MRHI, BCUT2D_MRLOW, AvgIpc, BalabanJ, HallKierAlpha, Ipc, PEOE_VSA7, SMR_VSA10, SMR_VSA7, SlogP_VSA2, TPSA, VSA_EState2, VSA_EState6, MolLogP, CalcAsphericity, CalcEccentricity, CalcInertialShapeFactor, CalcNPR2, CalcPBF, CalcPMI2, CalcSpherocityIndex | 36 |
| 44 | fluor | 2D | 1 | MaxAbsEStateIndex, MaxEStateIndex, MinAbsEStateIndex, MinEStateIndex, qed, SPS, MolWt, HeavyAtomMolWt, ExactMolWt, MaxPartialCharge, MinPartialCharge, MaxAbsPartialCharge, MinAbsPartialCharge, FpDensityMorgan1, FpDensityMorgan2, FpDensityMorgan3, BCUT2D_MWHI, BCUT2D_MWLOW, BCUT2D_CHGHI, BCUT2D_CHGLO, BCUT2D_LOGPHI, BCUT2D_LOGPLOW, BCUT2D_MRHI, BCUT2D_MRLOW, AvgIpc, BalabanJ, BertzCT, Chi0, Chi0n, Chi0v, Chi1, Chi1n, Chi1v, Chi2n, Chi2v, Chi3n, Chi3v, Chi4n, Chi4v, HallKierAlpha, Ipc, Kappa1, Kappa2, Kappa3, LabuteASA, PEOE_VSA7, SMR_VSA10, SMR_VSA7, SlogP_VSA2, SlogP_VSA6, TPSA, VSA_EState2, VSA_EState6, Phi, MolLogP, MolMR | 56 |

Continued on next page

| Cond. | Meas. | Dim. | Thresh. | Selected descriptors | ND |
|---|---|---|---|---|---|
| 45 | fluor | 3D | 1 | MaxAbsEStateIndex, MaxEStateIndex, MinAbsEStateIndex, MinEStateIndex, qed, SPS, MolWt, HeavyAtomMolWt, ExactMolWt, MaxPartialCharge, MinPartialCharge, MaxAbsPartialCharge, MinAbsPartialCharge, FpDensityMorgan1, FpDensityMorgan2, FpDensityMorgan3, BCUT2D_MWHI, BCUT2D_MWLOW, BCUT2D_CHGHI, BCUT2D_CHGLO, BCUT2D_LOGPHI, BCUT2D_LOGPLOW, BCUT2D_MRHI, BCUT2D_MRLOW, AvgIpc, BalabanJ, BertzCT, Chi0, Chi0n, Chi0v, Chi1, Chi1n, Chi1v, Chi2n, Chi2v, Chi3n, Chi3v, Chi4n, Chi4v, HallKierAlpha, Ipc, Kappa1, Kappa2, Kappa3, LabuteASA, PEOE_VSA7, SMR_VSA10, SMR_VSA7, SlogP_VSA2, SlogP_VSA6, TPSA, VSA_EState2, VSA_EState6, Phi, MolLogP, MolMR, CalcAsphericity, CalcChi0n, CalcChi0v, CalcChi1n, CalcChi1v, CalcChi2n, CalcChi2v, CalcChi3n, CalcChi3v, CalcChi4n, CalcChi4v, CalcEccentricity, CalcExactMolWt, CalcHallKierAlpha, CalcInertialShapeFactor, CalcKappa1, CalcKappa2, CalcKappa3, CalcLabuteASA, CalcNPR1, CalcNPR2, CalcPBF, CalcPMI1, CalcPMI2, CalcPMI3, CalcPhi, CalcRadiusOfGyration, CalcSpherocityIndex, CalcTPSA | 85 |

**Table S2. Molecular descriptors used in construction of SVM (model 9) for $M_{pro}$ inhibitors.** The table presents the molecular descriptors calculated for six main protease inhibitors with activity against SARS-CoV-2 (preclinical or clinical trials). Protonation states were assigned at pH 7.0 using a customized version of Dimorphite-DL, and all descriptors were subsequently computed using RDKit modules. In this table, 36 molecular descriptors used in construction of the model 9 (Support Vector Machine) were computated for boceprevir, ensitrelvir, lufotrelvir, nirmatrelvir, ritonavir, and simnotrelvir.

| Molecular descriptor | boceprevir | ensitrelvir | lufotrelvir | nirmatrelvir | ritonavir | simnotrelvir |
|---|---|---|---|---|---|---|
| MaxAbsEStateIndex | 13.91 | 14.49 | 13.26 | 13.47 | 13.87 | 13.61 |
| MinAbsEStateIndex | 0.08 | 0.15 | 0.03 | 0.04 | 0.07 | 0.12 |
| MinEStateIndex | -1.07 | -1.38 | -5.43 | -5.16 | -1.05 | -5.17 |
| qed | 0.36 | 0.33 | 0.24 | 0.50 | 0.11 | 0.46 |
| SPS | 26.27 | 11.41 | 17.21 | 28.89 | 13.68 | 25.25 |
| MolWt | 519.69 | 531.89 | 550.51 | 499.53 | 720.96 | 549.64 |
| MaxPartialCharge | 0.32 | 0.35 | 0.27 | 0.47 | 0.41 | 0.47 |
| MinPartialCharge | -0.36 | -0.32 | -0.79 | -0.36 | -0.44 | -0.36 |
| FpDensityMorgan1 | 1.00 | 0.92 | 1.18 | 1.23 | 0.90 | 1.22 |
| BCUT2D_MWHI | 16.18 | 35.50 | 31.20 | 19.41 | 32.13 | 32.22 |
| BCUT2D_MWLOW | 9.80 | 10.17 | 9.95 | 9.85 | 9.93 | 9.85 |
| BCUT2D_CHGHI | 2.70 | 2.19 | 2.33 | 2.70 | 2.31 | 2.54 |
| BCUT2D_CHGLO | -2.38 | -2.16 | -2.30 | -2.37 | -2.30 | -2.35 |
| BCUT2D_LOGPHI | 2.65 | 2.29 | 2.27 | 2.65 | 2.20 | 2.52 |
| BCUT2D_LOGPLOW | -2.62 | -2.43 | -2.53 | -2.61 | -2.55 | -2.59 |
| BCUT2D_MRHI | 6.37 | 6.34 | 7.43 | 5.95 | 7.09 | 8.21 |
| BCUT2D_MRLOW | -0.14 | 0.49 | -0.34 | -0.18 | -0.12 | -0.18 |
| AvgIpc | 3.06 | 3.40 | 3.46 | 3.10 | 3.79 | 3.50 |
| BalabanJ | 1.83 | 1.64 | 1.78 | 1.81 | 1.46 | 1.82 |
| HallKierAlpha | -2.65 | -4.46 | -3.25 | -2.84 | -4.33 | -2.14 |
| Ipc | 35028707.22 | 206483752.77 | 171410280.37 | 21524854.81 | 99661659139.94 | 53978979.66 |
| PEOE_VSA7 | 55.78 | 18.20 | 43.38 | 35.51 | 36.31 | 18.26 |
| SMR_VSA10 | 29.54 | 34.14 | 42.23 | 23.63 | 40.71 | 47.15 |
| SMR_VSA7 | 0.00 | 91.62 | 29.96 | 0.00 | 104.46 | 0.00 |
| SlogP_VSA2 | 64.65 | 38.66 | 60.83 | 65.92 | 69.28 | 81.50 |
| TPSA | 150.70 | 117.45 | 201.81 | 131.40 | 145.78 | 131.40 |
| VSA_EState2 | 66.09 | 34.22 | 75.77 | 51.58 | 50.98 | 51.81 |
| VSA_EState6 | -3.13 | 4.20 | 4.50 | -1.48 | 16.79 | -1.53 |
| MolLogP | 1.71 | 2.33 | -0.25 | 1.10 | 5.91 | 1.39 |
| CalcAsphericity | 0.34 | 0.13 | 0.38 | 0.43 | 0.27 | 0.24 |
| CalcEccentricity | 0.95 | 0.82 | 0.96 | 0.97 | 0.93 | 0.92 |
| CalcInertialShapeFactor | 0.00 | 0.00 | 0.00 | 0.00 | 0.00 | 0.00 |
| CalcNPR2 | 0.92 | 0.63 | 0.81 | 0.85 | 0.81 | 0.69 |
| CalcPBF | 1.46 | 1.22 | 1.11 | 1.01 | 1.50 | 1.04 |
| CalcPMI2 | 10701.72 | 6555.23 | 12136.46 | 9588.80 | 17120.64 | 8164.86 |
| CalcSpherocityIndex | 0.33 | 0.26 | 0.18 | 0.19 | 0.30 | 0.23 |

**Table S3. Molecular descriptors used in construction of LR (model 10) for $M_{pro}$ inhibitors.** The table presents the molecular descriptors calculated for six main protease inhibitors with activity against SARS-CoV-2 (preclinical or clinical trials). Protonation states were assigned at pH 7.0 using a customized version of Dimorphite-DL, and all descriptors were subsequently computed using RDKit modules. In this table, 85 molecular descriptors used in construction of the model 10 (logistic regression) were computated for boceprevir, ensitrelvir, lufotrelvir, nirmatrelvir, ritonavir, and simnotrelvir.

| Molecular descriptor | boceprevir | ensitrelvir | lufotrelvir | nirmatrelvir | ritonavir | simnotrelvir |
|---|---|---|---|---|---|---|
| MaxAbsEStateIndex | 13.91 | 14.49 | 13.26 | 13.47 | 13.87 | 13.61 |
| MaxEStateIndex | 13.91 | 14.49 | 13.26 | 13.47 | 13.87 | 13.61 |
| MinAbsEStateIndex | 0.08 | 0.15 | 0.03 | 0.04 | 0.07 | 0.12 |
| MinEStateIndex | -1.07 | -1.38 | -5.43 | -5.16 | -1.05 | -5.17 |
| qed | 0.36 | 0.33 | 0.24 | 0.50 | 0.11 | 0.46 |
| SPS | 26.27 | 11.41 | 17.21 | 28.89 | 13.68 | 25.25 |
| MolWt | 519.69 | 531.89 | 550.51 | 499.53 | 720.96 | 549.64 |
| HeavyAtomMolWt | 474.33 | 514.75 | 519.26 | 467.28 | 672.58 | 519.40 |
| ExactMolWt | 519.34 | 531.11 | 550.18 | 499.24 | 720.31 | 549.17 |
| MaxPartialCharge | 0.32 | 0.35 | 0.27 | 0.47 | 0.41 | 0.47 |
| MinPartialCharge | -0.36 | -0.32 | -0.79 | -0.36 | -0.44 | -0.36 |
| MaxAbsPartialCharge | 0.36 | 0.35 | 0.79 | 0.47 | 0.44 | 0.47 |
| MinAbsPartialCharge | 0.32 | 0.32 | 0.27 | 0.36 | 0.41 | 0.36 |
| FpDensityMorgan1 | 1.00 | 0.92 | 1.18 | 1.23 | 0.90 | 1.22 |
| FpDensityMorgan2 | 1.59 | 1.68 | 1.92 | 1.86 | 1.56 | 1.86 |
| FpDensityMorgan3 | 2.05 | 2.41 | 2.53 | 2.34 | 2.14 | 2.36 |
| BCUT2D_MWHI | 16.18 | 35.50 | 31.20 | 19.41 | 32.13 | 32.22 |
| BCUT2D_MWLOW | 9.80 | 10.17 | 9.95 | 9.85 | 9.93 | 9.85 |
| BCUT2D_CHGHI | 2.70 | 2.19 | 2.33 | 2.70 | 2.31 | 2.54 |
| BCUT2D_CHGLO | -2.38 | -2.16 | -2.30 | -2.37 | -2.30 | -2.35 |
| BCUT2D_LOGPHI | 2.65 | 2.29 | 2.27 | 2.65 | 2.20 | 2.52 |
| BCUT2D_LOGPLOW | -2.62 | -2.43 | -2.53 | -2.61 | -2.55 | -2.59 |
| BCUT2D_MRHI | 6.37 | 6.34 | 7.43 | 5.95 | 7.09 | 8.21 |
| BCUT2D_MRLOW | -0.14 | 0.49 | -0.34 | -0.18 | -0.12 | -0.18 |
| AvgIpc | 3.06 | 3.40 | 3.46 | 3.10 | 3.79 | 3.50 |
| BalabanJ | 1.83 | 1.64 | 1.78 | 1.81 | 1.46 | 1.82 |
| BertzCT | 959.21 | 1791.04 | 1238.70 | 952.27 | 1617.50 | 943.44 |
| Chi0 | 28.09 | 26.43 | 28.03 | 26.51 | 36.05 | 26.89 |
| Chi0n | 23.57 | 19.28 | 21.28 | 20.45 | 29.24 | 20.24 |
| Chi0v | 23.57 | 20.04 | 22.17 | 20.45 | 30.88 | 21.87 |
| Chi1 | 16.84 | 17.58 | 17.87 | 16.00 | 23.98 | 16.67 |
| Chi1n | 13.37 | 10.68 | 12.12 | 11.66 | 16.81 | 11.61 |
| Chi1v | 13.37 | 11.06 | 13.58 | 11.66 | 18.57 | 13.58 |
| Chi2n | 13.86 | 8.18 | 9.52 | 11.36 | 13.04 | 10.27 |
| Chi2v | 13.86 | 8.59 | 10.67 | 11.36 | 14.90 | 13.48 |
| Chi3n | 8.31 | 5.47 | 6.17 | 7.56 | 8.13 | 6.49 |
| Chi3v | 8.31 | 5.78 | 6.62 | 7.56 | 10.08 | 9.92 |
| Chi4n | 5.86 | 3.74 | 4.31 | 5.40 | 5.39 | 4.60 |
| Chi4v | 5.86 | 3.95 | 4.59 | 5.40 | 6.72 | 7.86 |
| HallKierAlpha | -2.65 | -4.46 | -3.25 | -2.84 | -4.33 | -2.14 |
| Ipc | 35028707.22 | 206483752.77 | 171410280.37 | 21524854.81 | 99661659139.94 | 53978979.66 |
| Kappa1 | 28.91 | 24.24 | 29.31 | 26.76 | 38.47 | 28.43 |
| Kappa2 | 9.91 | 9.20 | 12.53 | 8.99 | 19.17 | 10.69 |
| Kappa3 | 6.26 | 4.70 | 7.87 | 5.23 | 12.22 | 6.40 |
| LabuteASA | 219.77 | 209.93 | 218.82 | 201.70 | 302.06 | 215.58 |

Continued on next page

| Molecular descriptor | boceprevir | ensitrelvir | lufotrelvir | nirmatrelvir | ritonavir | simnotrelvir |
|---|---|---|---|---|---|---|
| PEOE_VSA7 | 55.78 | 18.20 | 43.38 | 35.51 | 36.31 | 18.26 |
| SMR_VSA10 | 29.54 | 34.14 | 42.23 | 23.63 | 40.71 | 47.15 |
| SMR_VSA7 | 0.00 | 91.62 | 29.96 | 0.00 | 104.46 | 0.00 |
| SlogP_VSA2 | 64.65 | 38.66 | 60.83 | 65.92 | 69.28 | 81.50 |
| SlogP_VSA6 | 0.00 | 46.38 | 24.27 | 0.00 | 77.75 | 0.00 |
| TPSA | 150.70 | 117.45 | 201.81 | 131.40 | 145.78 | 131.40 |
| VSA_EState2 | 66.09 | 34.22 | 75.77 | 51.58 | 50.98 | 51.81 |
| VSA_EState6 | -3.13 | 4.20 | 4.50 | -1.48 | 16.79 | -1.53 |
| Phi | 7.74 | 6.03 | 9.66 | 6.87 | 14.75 | 8.44 |
| MolLogP | 1.71 | 2.33 | -0.25 | 1.10 | 5.91 | 1.39 |
| MolMR | 139.02 | 127.29 | 131.79 | 116.98 | 196.90 | 128.48 |
| CalcAsphericity | 0.34 | 0.13 | 0.38 | 0.43 | 0.27 | 0.24 |
| CalcChi0n | 62.78 | 34.35 | 48.91 | 48.50 | 72.04 | 46.82 |
| CalcChi0v | 17.78 | 18.11 | 18.80 | 16.50 | 25.67 | 18.45 |
| CalcChi1n | 31.49 | 17.79 | 24.49 | 24.46 | 36.20 | 23.53 |
| CalcChi1v | 9.26 | 9.72 | 10.66 | 8.62 | 14.08 | 10.32 |
| CalcChi2n | 7.19 | 6.66 | 6.27 | 6.65 | 8.06 | 6.30 |
| CalcChi2v | 7.19 | 7.04 | 7.35 | 6.65 | 9.65 | 8.59 |
| CalcChi3n | 4.30 | 4.26 | 3.73 | 4.16 | 4.66 | 3.73 |
| CalcChi3v | 4.30 | 4.54 | 4.04 | 4.16 | 6.00 | 5.73 |
| CalcChi4n | 2.71 | 2.75 | 2.30 | 2.60 | 2.73 | 2.29 |
| CalcChi4v | 2.71 | 2.93 | 2.50 | 2.60 | 3.61 | 3.90 |
| CalcEccentricity | 0.95 | 0.82 | 0.96 | 0.97 | 0.93 | 0.92 |
| CalcExactMolWt | 519.34 | 531.11 | 550.18 | 499.24 | 720.31 | 549.17 |
| CalcHallKierAlpha | -2.65 | -4.46 | -3.25 | -2.84 | -4.33 | -2.14 |
| CalcInertialShapeFactor | 0.00 | 0.00 | 0.00 | 0.00 | 0.00 | 0.00 |
| CalcKappa1 | 5.77 | 11.29 | 8.62 | 7.13 | 9.75 | 8.43 |
| CalcKappa2 | 9.91 | 9.20 | 12.53 | 8.99 | 19.17 | 10.69 |
| CalcKappa3 | 6.26 | 4.70 | 7.87 | 5.23 | 12.22 | 6.40 |
| CalcLabuteASA | 283.28 | 233.87 | 262.60 | 246.83 | 369.76 | 257.90 |
| CalcNPR1 | 0.31 | 0.57 | 0.28 | 0.25 | 0.37 | 0.40 |
| CalcNPR2 | 0.92 | 0.63 | 0.81 | 0.85 | 0.81 | 0.69 |
| CalcPBF | 1.46 | 1.22 | 1.11 | 1.01 | 1.50 | 1.04 |
| CalcPMI1 | 3604.53 | 5925.62 | 4264.20 | 2773.28 | 7744.19 | 4738.20 |
| CalcPMI2 | 10701.72 | 6555.23 | 12136.46 | 9588.80 | 17120.64 | 8164.86 |
| CalcPMI3 | 11664.18 | 10326.55 | 14981.61 | 11254.41 | 21123.99 | 11772.97 |
| CalcPhi | 1.55 | 2.81 | 2.84 | 1.83 | 3.74 | 2.50 |
| CalcRadiusOfGyration | 5.00 | 4.63 | 5.34 | 4.86 | 5.65 | 4.74 |
| CalcSpherocityIndex | 0.33 | 0.26 | 0.18 | 0.19 | 0.30 | 0.23 |
| CalcTPSA | 150.70 | 117.45 | 201.81 | 131.40 | 145.78 | 131.40 |

**Table S4. Statistical summary of molecular descriptors for the $M_{pro}$ inhibitors in preclinical or clinical trials.** The table shows the 95% confidence interval (IC_95) (inferior and superior), mean, and standard deviation (SD) for 36 molecular descriptors used in SVM (model 9) calculated for the $M_{pro}$ inhibitors in preclinical or clinical trials.

| Molecular descriptor | IC95_inf | IC95_sup | Mean | SD |
|---|---|---|---|---|
| MaxAbsEStateIndex | 13.32 | 14.22 | 13.77 | 0.43 |
| MinAbsEStateIndex | 0.03 | 0.13 | 0.08 | 0.05 |
| MinEStateIndex | -5.56 | -0.86 | -3.21 | 2.24 |
| qed | 0.18 | 0.48 | 0.33 | 0.14 |
| SPS | 12.79 | 28.11 | 20.45 | 7.3 |
| MolWt | 477.88 | 646.19 | 562.04 | 80.19 |
| MaxPartialCharge | 0.3 | 0.47 | 0.38 | 0.08 |
| MinPartialCharge | -0.62 | -0.25 | -0.44 | 0.18 |
| FpDensityMorgan1 | 0.91 | 1.24 | 1.07 | 0.15 |
| BCUT2D_MWHI | 19.45 | 36.1 | 27.77 | 7.93 |
| BCUT2D_MWLOW | 9.79 | 10.06 | 9.92 | 0.13 |
| BCUT2D_CHGHI | 2.23 | 2.69 | 2.46 | 0.22 |
| BCUT2D_CHGLO | -2.39 | -2.23 | -2.31 | 0.08 |
| BCUT2D_LOGPHI | 2.22 | 2.64 | 2.43 | 0.2 |
| BCUT2D_LOGPLOW | -2.63 | -2.48 | -2.55 | 0.07 |
| BCUT2D_MRHI | 6.02 | 7.78 | 6.9 | 0.84 |
| BCUT2D_MRLOW | -0.38 | 0.22 | -0.08 | 0.29 |
| AvgIpc | 3.1 | 3.67 | 3.38 | 0.27 |
| BalabanJ | 1.57 | 1.88 | 1.72 | 0.15 |
| HallKierAlpha | -4.26 | -2.3 | -3.28 | 0.94 |
| Ipc | -25964617468.43 | 59347979373.35 | 16691680952.46 | 40646893228.14 |
| PEOE_VSA7 | 19.25 | 49.89 | 34.57 | 14.6 |
| SMR_VSA10 | 27.05 | 45.42 | 36.23 | 8.75 |
| SMR_VSA7 | -13.07 | 88.41 | 37.67 | 48.35 |
| SlogP_VSA2 | 48.72 | 78.23 | 63.47 | 14.06 |
| TPSA | 115.37 | 177.47 | 146.42 | 29.59 |
| VSA_EState2 | 40.06 | 70.09 | 55.07 | 14.31 |
| VSA_EState6 | -4.51 | 10.96 | 3.22 | 7.37 |
| MolLogP | -0.16 | 4.22 | 2.03 | 2.08 |
| CalcAsphericity | 0.19 | 0.41 | 0.3 | 0.11 |
| CalcEccentricity | 0.87 | 0.98 | 0.92 | 0.05 |
| CalcInertialShapeFactor | 0 | 0 | 0 | 0 |
| CalcNPR2 | 0.67 | 0.9 | 0.79 | 0.11 |
| CalcPBF | 1 | 1.45 | 1.22 | 0.21 |
| CalcPMI2 | 6837.51 | 14585.06 | 10711.28 | 3691.29 |
| CalcSpherocityIndex | 0.19 | 0.31 | 0.25 | 0.06 |

**Table S5. Statistical summary of molecular descriptors for the $M_{pro}$ inhibitors in preclinical or clinical trials.** The table shows the 95% confidence interval (IC_95) (inferior and superior), mean, and standard deviation (SD) for 85 molecular descriptors used in LR (model 10) calculated for the $M_{pro}$ inhibitors in preclinical or clinical trials.

| Molecular descriptor | IC95_inf | IC95_sup | Mean | SD |
|---|---|---|---|---|
| MaxAbsEStateIndex | 13.32 | 14.22 | 13.77 | 0.43 |
| MaxEStateIndex | 13.32 | 14.22 | 13.77 | 0.43 |
| MinAbsEStateIndex | 0.03 | 0.13 | 0.08 | 0.05 |
| MinEStateIndex | -5.56 | -0.86 | -3.21 | 2.24 |
| qed | 0.18 | 0.48 | 0.33 | 0.14 |
| SPS | 12.79 | 28.11 | 20.45 | 7.3 |
| MolWt | 477.88 | 646.19 | 562.04 | 80.19 |
| HeavyAtomMolWt | 449.69 | 606.18 | 527.93 | 74.56 |
| ExactMolWt | 477.5 | 645.62 | 561.56 | 80.1 |
| MaxPartialCharge | 0.3 | 0.47 | 0.38 | 0.08 |
| MinPartialCharge | -0.62 | -0.25 | -0.44 | 0.18 |
| MaxAbsPartialCharge | 0.31 | 0.65 | 0.48 | 0.16 |
| MinAbsPartialCharge | 0.29 | 0.39 | 0.34 | 0.05 |
| FpDensityMorgan1 | 0.91 | 1.24 | 1.07 | 0.15 |
| FpDensityMorgan2 | 1.58 | 1.91 | 1.74 | 0.15 |
| FpDensityMorgan3 | 2.12 | 2.49 | 2.31 | 0.18 |
| BCUT2D_MWHI | 19.45 | 36.1 | 27.77 | 7.93 |
| BCUT2D_MWLOW | 9.79 | 10.06 | 9.92 | 0.13 |
| BCUT2D_CHGHI | 2.23 | 2.69 | 2.46 | 0.22 |
| BCUT2D_CHGLO | -2.39 | -2.23 | -2.31 | 0.08 |
| BCUT2D_LOGPHI | 2.22 | 2.64 | 2.43 | 0.2 |
| BCUT2D_LOGPLOW | -2.63 | -2.48 | -2.55 | 0.07 |
| BCUT2D_MRHI | 6.02 | 7.78 | 6.9 | 0.84 |
| BCUT2D_MRLOW | -0.38 | 0.22 | -0.08 | 0.29 |
| AvgIpc | 3.1 | 3.67 | 3.38 | 0.27 |
| BalabanJ | 1.57 | 1.88 | 1.72 | 0.15 |
| BertzCT | 859.08 | 1641.64 | 1250.36 | 372.85 |
| Chi0 | 24.79 | 32.54 | 28.67 | 3.69 |
| Chi0n | 18.48 | 26.2 | 22.34 | 3.68 |
| Chi0v | 18.98 | 27.35 | 23.16 | 3.99 |
| Chi1 | 15.08 | 21.23 | 18.16 | 2.93 |
| Chi1n | 10.41 | 15.01 | 12.71 | 2.19 |
| Chi1v | 10.86 | 16.41 | 13.64 | 2.64 |
| Chi2n | 8.78 | 13.3 | 11.04 | 2.15 |
| Chi2v | 9.67 | 14.62 | 12.14 | 2.36 |
| Chi3n | 5.82 | 8.23 | 7.02 | 1.15 |
| Chi3v | 6.22 | 9.87 | 8.04 | 1.74 |
| Chi4n | 4.04 | 5.72 | 4.88 | 0.8 |
| Chi4v | 4.24 | 7.22 | 5.73 | 1.42 |
| HallKierAlpha | -4.26 | -2.3 | -3.28 | 0.94 |
| Ipc | -25964617468.43 | 59347979373.35 | 16691680952.46 | 40646893228.14 |
| Kappa1 | 24.28 | 34.43 | 29.35 | 4.84 |
| Kappa2 | 7.7 | 15.79 | 11.75 | 3.85 |
| Kappa3 | 4.25 | 9.98 | 7.11 | 2.73 |
| LabuteASA | 189.25 | 266.7 | 227.98 | 36.9 |
| PEOE_VSA7 | 19.25 | 49.89 | 34.57 | 14.6 |
| SMR_VSA10 | 27.05 | 45.42 | 36.23 | 8.75 |
| SMR_VSA7 | -13.07 | 88.41 | 37.67 | 48.35 |

Continued on next page

| Molecular descriptor | IC95_inf | IC95_sup | Mean | SD |
|---|---|---|---|---|
| SlogP_VSA2 | 48.72 | 78.23 | 63.47 | 14.06 |
| SlogP_VSA6 | -8.83 | 58.3 | 24.73 | 31.98 |
| TPSA | 115.37 | 177.47 | 146.42 | 29.59 |
| VSA_EState2 | 40.06 | 70.09 | 55.07 | 14.31 |
| VSA_EState6 | -4.51 | 10.96 | 3.22 | 7.37 |
| Phi | 5.64 | 12.19 | 8.91 | 3.12 |
| MolLogP | -0.16 | 4.22 | 2.03 | 2.08 |
| MolMR | 109.92 | 170.24 | 140.08 | 28.74 |
| CalcAsphericity | 0.19 | 0.41 | 0.3 | 0.11 |
| CalcChi0n | 38.33 | 66.14 | 52.23 | 13.25 |
| CalcChi0v | 15.8 | 22.64 | 19.22 | 3.26 |
| CalcChi1n | 19.5 | 33.16 | 26.33 | 6.51 |
| CalcChi1v | 8.42 | 12.46 | 10.44 | 1.93 |
| CalcChi2n | 6.14 | 7.57 | 6.86 | 0.68 |
| CalcChi2v | 6.55 | 8.94 | 7.75 | 1.14 |
| CalcChi3n | 3.76 | 4.52 | 4.14 | 0.36 |
| CalcChi3v | 3.9 | 5.69 | 4.79 | 0.85 |
| CalcChi4n | 2.34 | 2.79 | 2.56 | 0.21 |
| CalcChi4v | 2.44 | 3.65 | 3.04 | 0.58 |
| CalcEccentricity | 0.87 | 0.98 | 0.92 | 0.05 |
| CalcExactMolWt | 477.5 | 645.62 | 561.56 | 80.1 |
| CalcHallKierAlpha | -4.26 | -2.3 | -3.28 | 0.94 |
| CalcInertialShapeFactor | 0 | 0 | 0 | 0 |
| CalcKappa1 | 6.47 | 10.53 | 8.5 | 1.93 |
| CalcKappa2 | 7.7 | 15.79 | 11.75 | 3.85 |
| CalcKappa3 | 4.25 | 9.98 | 7.11 | 2.73 |
| CalcLabuteASA | 224.35 | 327.07 | 275.71 | 48.94 |
| CalcNPR1 | 0.24 | 0.48 | 0.36 | 0.12 |
| CalcNPR2 | 0.67 | 0.9 | 0.79 | 0.11 |
| CalcPBF | 1 | 1.45 | 1.22 | 0.21 |
| CalcPMI1 | 2978.96 | 6704.38 | 4841.67 | 1774.96 |
| CalcPMI2 | 6837.51 | 14585.06 | 10711.28 | 3691.29 |
| CalcPMI3 | 9276.32 | 17764.91 | 13520.62 | 4044.36 |
| CalcPhi | 1.72 | 3.37 | 2.55 | 0.79 |
| CalcRadiusOfGyration | 4.63 | 5.44 | 5.04 | 0.39 |
| CalcSpherocityIndex | 0.19 | 0.31 | 0.25 | 0.06 |
| CalcTPSA | 115.37 | 177.47 | 146.42 | 29.59 |

### Figures

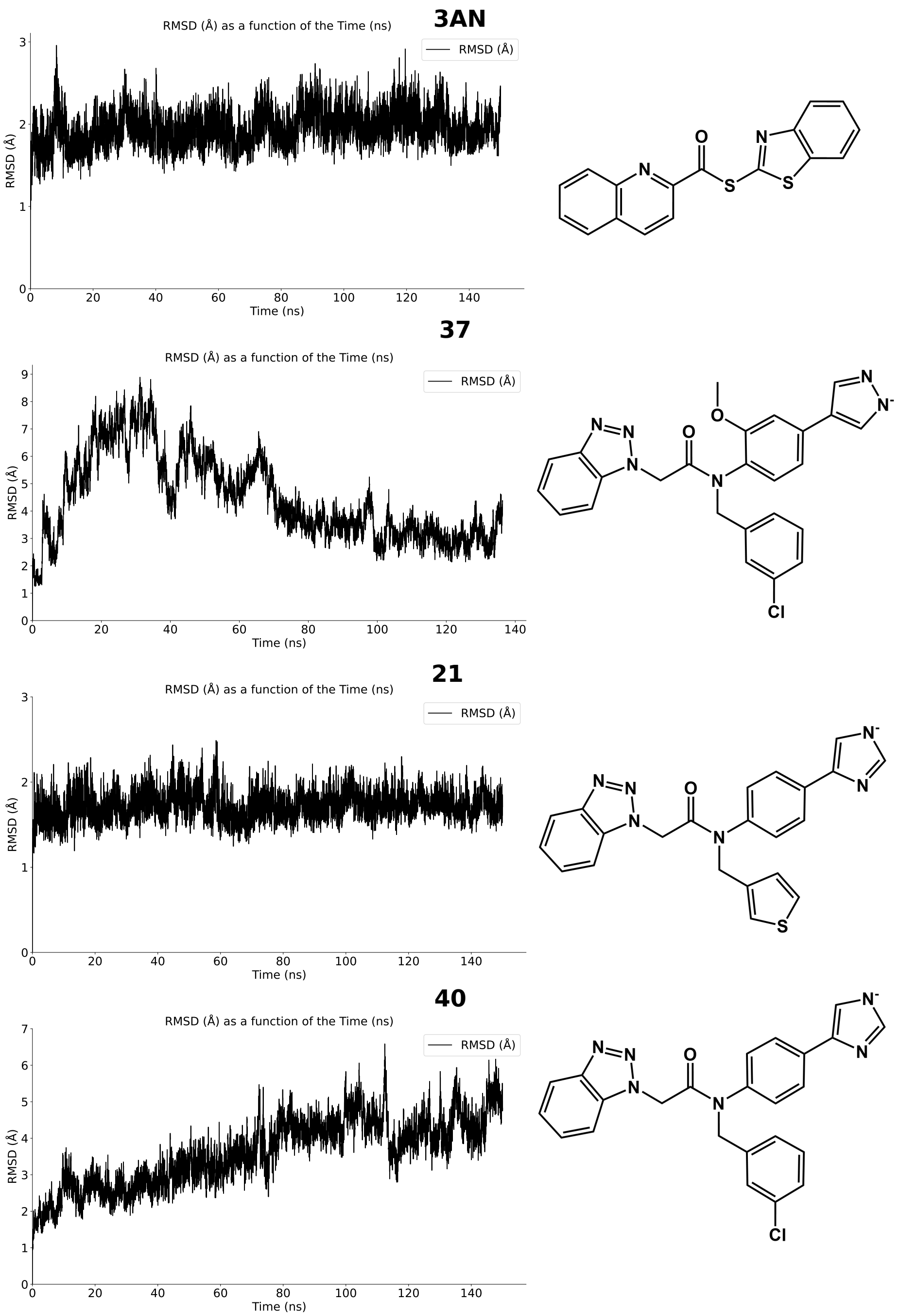

**Figure S1. Backbone root-mean-square deviation (RMSD) of $M_{pro}$ in complex with different inhibitors.** Backbone RMSD values were calculated from molecular dynamics (MD) simulations of $M_{pro}$ bound to inhibitors 3AN, 37, 21, and 40.

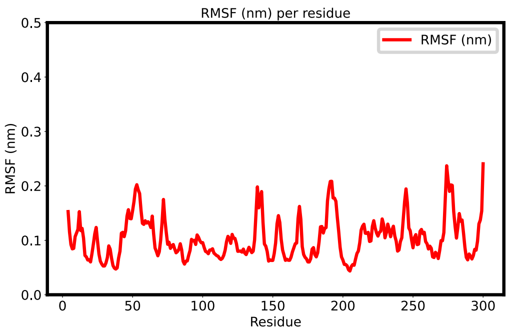

**3AN**

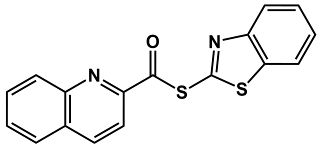

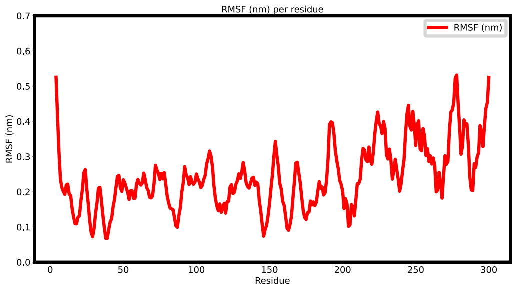

**37**

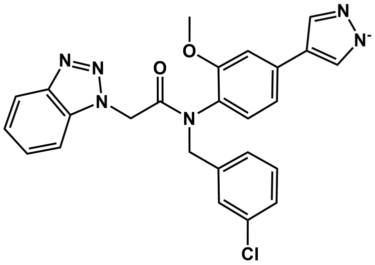

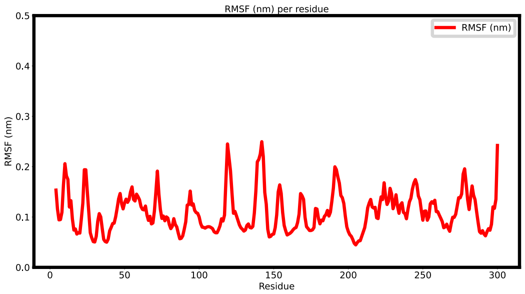

**21**

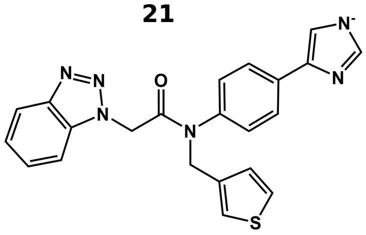

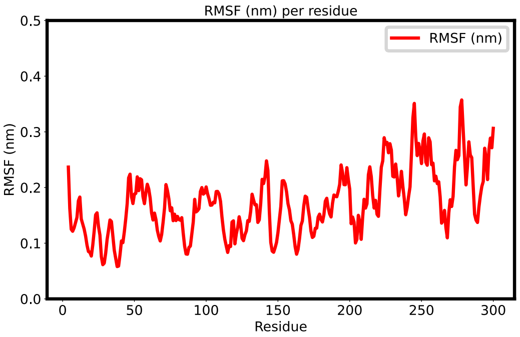

**40**

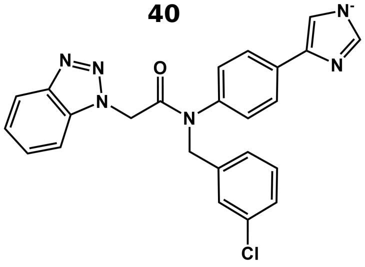

**Figure S2. Backbone root-mean-square fluctuation (RMSF) of $M_{pro}$ in complex with different inhibitors.** Backbone RMSF values were obtained from MD trajectories of $M_{pro}$ in complex with inhibitors 3AN, 37, 21, and 40.

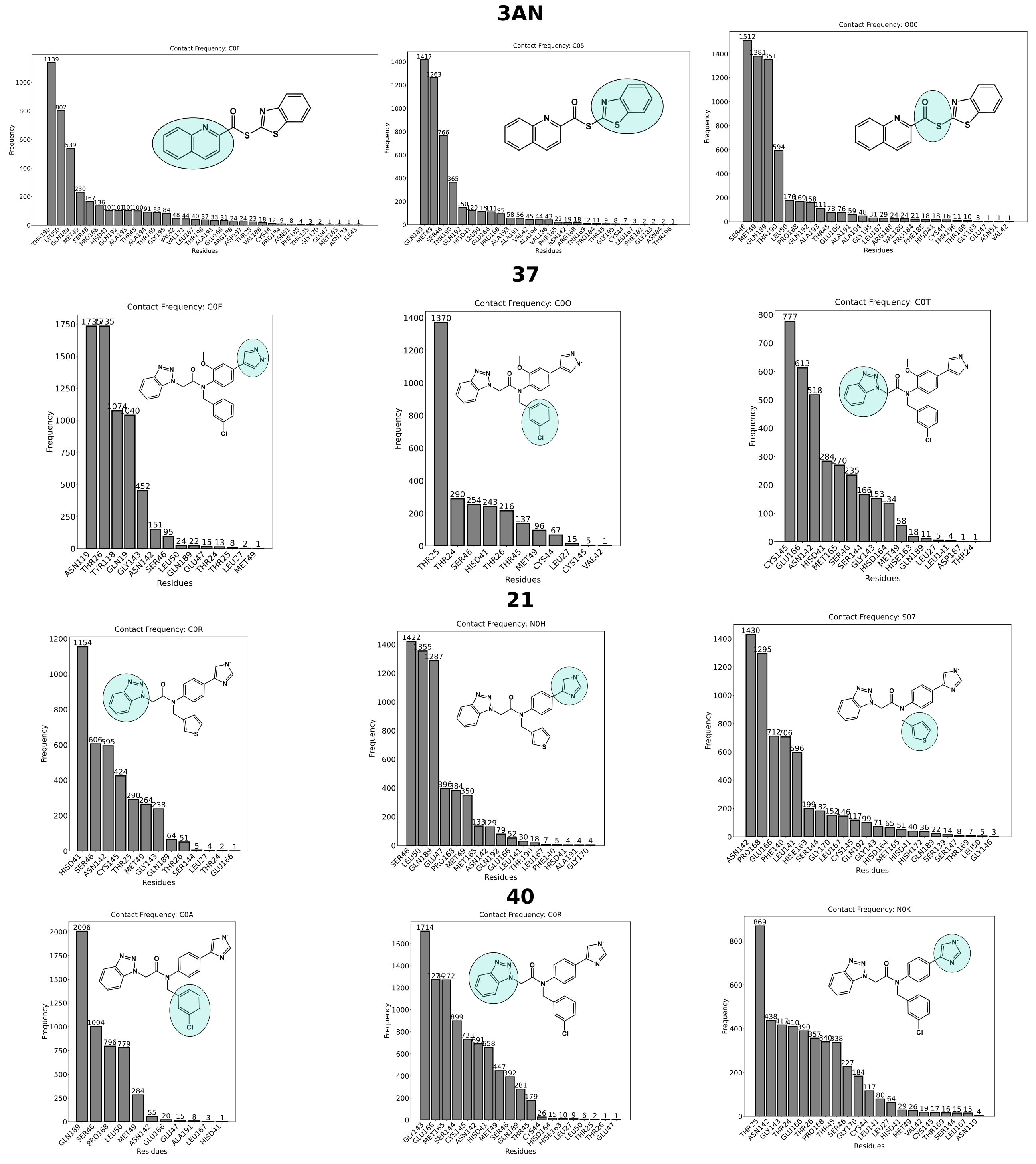

**Figure S3. Contacts of chemical groups of inhibitors in active site of SARS-CoV-2 M$^{pro}$.** The contact analysis considered three distant atoms (and different chemical groups) (blue ellipses) that compose the inhibitor: 3AN [C0F (quinoline), C05 (benzothiazole), O00 (thioamide)], 37 [C0F (pyrazole), C0O (chlorophenyl), C0T (benzotriazole)], 21 [C0R (benzotriazole), S07 (thiophene), N0H (imidazole)], and 40 [C0R (benzotriazole), N0K (imidazole), C0A (chlorophenyl)]. The coordinates of these atoms define the center of a sphere with a radius of 4 Å. All atoms within the sphere were considered potential residues for interaction and the subsite with which the inhibitor interacted was assigned. Next, the contact frequency per residue was calculated.
